# Overexpression-based detection of translatable circular RNAs is vulnerable to coexistent linear RNA byproducts

**DOI:** 10.1101/2021.03.23.433163

**Authors:** Yanyi Jiang, Xiaofan Chen, Wei Zhang

## Abstract

In RNA field, the demarcation between coding and non-coding has been negotiated by the recent discovery of occasionally translated circular RNAs (circRNAs). Although absent of 5’ cap structure, circRNAs can be translated cap-independently. Complementary intron-mediated overexpression is one of the most utilized methodologies for circRNA research but not without bearing echoing skepticism for its poorly defined mechanism and latent coexistent side products. In this study, leveraging such circRNA overexpression system, we have interrogated the protein-coding potential of 30 human circRNAs containing infinite open reading frames in HEK293T cells. Surprisingly, pervasive translation signals are detected by immunoblotting. However, intensive mutagenesis reveals that numerous translation signals are generated independently of circRNA synthesis. We have developed a dual tag strategy to isolate translation noise and directly demonstrate that the fallacious translation signals originate from cryptically spliced linear transcripts. The concomitant linear RNA byproducts, presumably concatemers, can be translated to allow pseudo rolling circle translation signals, and can involve backsplicing junction (BSJ) to disqualify the BSJ-based evidence for circRNA translation. We also find non-AUG start codons may engage in the translation initiation of circRNAs. Taken together, our systematic evaluation sheds light on heterogeneous translational outputs from circRNA overexpression vector and comes with a caveat that ectopic overexpression technique necessitates extremely rigorous control setup in circRNA translation and functional investigation.

## 1. Introduction

Circular RNAs (circRNAs), generated from backsplicing in which the splice donor of an exon covalently joints the splice acceptor site of an upstream exon, have been thrusted into the biomedicine limelight for their high abundance and versatile biological function [1-4]. CircRNAs have recently attracted accumulating attention for their protein-coding potential, even in the absence of 5’ cap and 3’ poly(A) tail structures that both sustain the canonical cap-dependent translation machinery. It has been reported that circRNAs can serve as templates for cap-independent translation via virus or cellular internal ribosome entry site (IRES) [5-7], m6A modification [8], short IRES-like sequence [9], and other elusive mechanisms [10]. While the translation of circRNAs and their cellular impact cannot reach consensus [11-16], tentative and massive efforts have been made to identify endogenous translatable circRNAs [7], investigate the function of endogenous circRNA-derived peptides [17,18], and explore the bioengineering application for protein production [19-22]. The circular structure renders circRNA not only tolerant to RNA exonuclease resulting in high stability [11,23], but also the possibility to enable rolling circle translation from infinite open reading frames (ORFs), ORFs without stop codons [5,9,10,19].

Among current approaches for circRNA study, the complementary intron-mediated circularization stands out as a preferred widely used methodology for circRNA overexpression, as flanking intronic complementary sequence efficiently facilitates backsplicing to spur circRNA biogenesis [24-26]. Unfortunately, coexistent linear overlapping sequence seems inevitable and usually entangles with other side products including concatenated RNAs [7,20,27,28]. These byproducts are prone to undermining experimental design and inducing misleading conclusions. For example, circ-ZNF609 was reported to be translationally active [29], but a following study has denied this possibility [28], as the putative overexpressed circRNA-derived peptides were even generated when the synthesis of circ-ZNF609 was compromised. Thus, the dark side of circRNA overexpression strategy deserves a closer inspection, whereas the methodical evaluation remains limited.

Here, using FLAG tagged ZKSCAN1 minigene circRNA overexpression vector [26], we have conducted a coding potential assessment of 30 human circRNAs carrying infinite ORFs and annotated AUG start codons. While pervasive translation events are detected by immunoblotting, following mutagenesis study demonstrates many translation signals are not from circRNAs. Such aberrant translation is initiated independently of the AUG start codons within the circRNA sequence but can be masked upon epitope tags spanning backsplicing junction (BSJ). A FLAG-Myc dual tag reporter is developed to discriminate translation noise of linear RNAs. Using this system, we ascertain that the fallacious translation events originate from concomitant cryptically spliced linear RNAs. Unexpectedly, we further determine the translation of BSJ-involved linear RNA byproducts and identify linear RNA-derived pseudo rolling circle translation signals. Additionally, we find non-AUG start codons may participate in the translation initiation of circRNAs.

## 2. Materials and methods

### 2.1. Plasmids

All the circular RNA overexpression constructs used pcDNA3.1(+) CircRNA Mini Vector-ZKSCAN1 (Addgene, Plasmid #60649) [26] as backbone. The splicing signals of all constructs were predetermined according to Human Splicing Finder [30] or “Splice Site Prediction by Neural Network” website NNSPLICE 0.9 version. To optimize splicing strength and make sure FLAG is in a right ORF, a subset of circRNA sequence was adjusted, including frame shifting and adding CAG codon upstream of splice donor. All constructs and mutants were created through standard PCR-based mutagenesis method and confirmed by DNA sequencing. Additional details and sequence of all constructs are provided in the Supplemental Materials (Table 1, Fig. S2, S3).

**Table 1.**
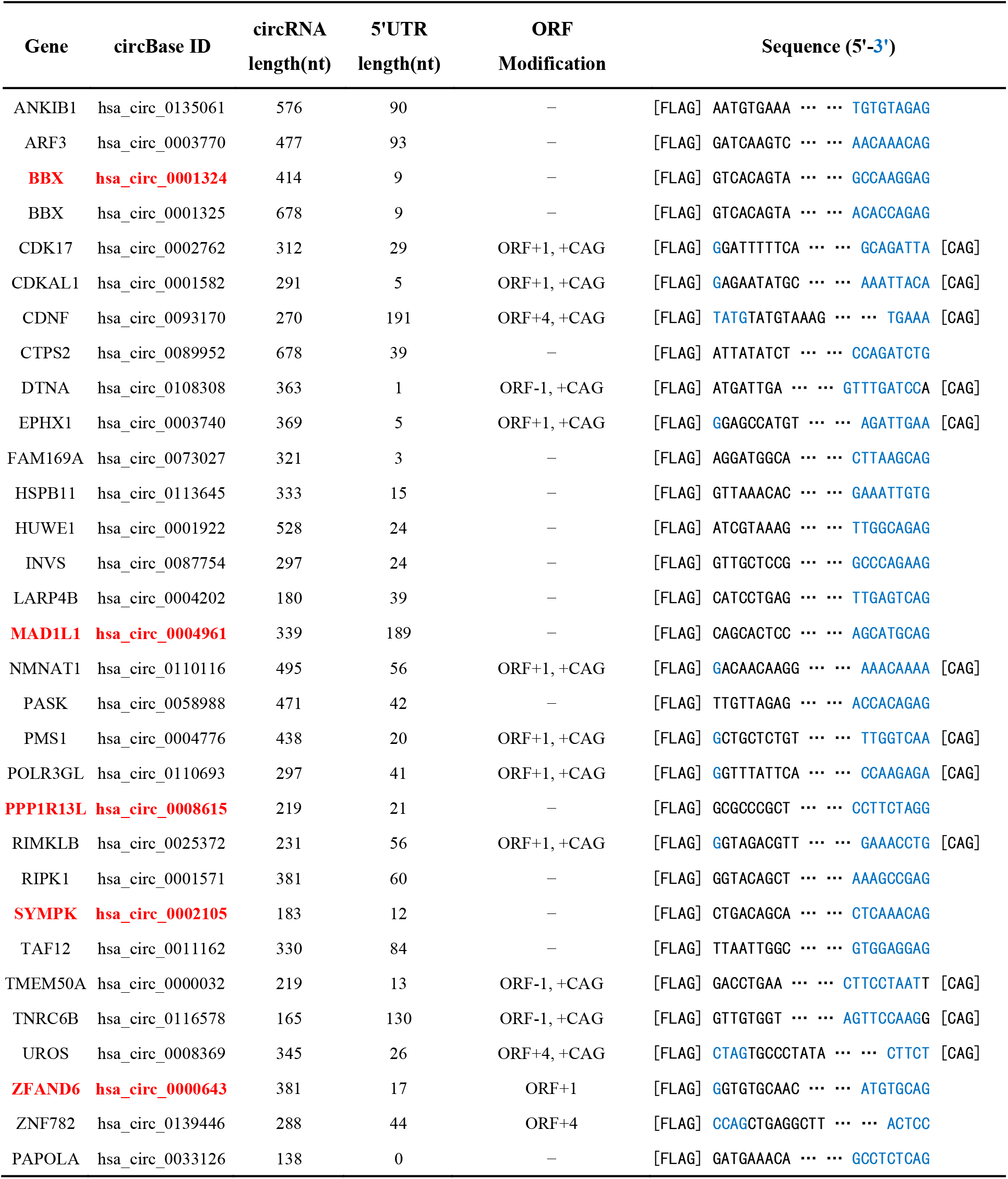
The circRNAs for coding potential investigation. Several ORFs of circRNA are adjusted (see methods). The sequence detail is described here. Five shortlisted circRNA candidates are highlighted in red. The annotated 3’ terminal of circRNA sequence is marked in blue.

### 2.2. Cell Culture and transfection

HEK293T (ATCC) were cultured in Dulbecco’s Modified Eagle Medium (DMEM) with 10% fetal bovine serum (Gibco), and 1% of penicillin-streptomycin at 37°C with 5% CO2. Plasmids were transfected into HEK293T with ViaFect (#E4982, Promega). 2.5ng of total DNA was added per well in a 6-well plate at reagent:DNA ratio of 4:1. Cells were harvested after 42-50h without medium change.

### 2.3. Western blotting

Total proteins from HEK293T cells were extracted using lysis buffer (150mM NaCl, 50 mM Tris-HCl, pH7.4, 0.5% Triton X-100) adding protease inhibitor cocktail (#11836153001, Roche). Prepared protein samples were separated by 12% or 15% SDS-PAGE and subsequently transferred onto a 0.2μm PVDF membrane (Pall). The immunoblotting was performed with the following primary antibodies (overnight at 4°C) and secondary antibodies (1h at RT): mouse-anti-Flag (M2 monoclonal, #F3165, 1:3000, Sigma), rabbit-anti-GAPDH (monoclonal, #2118, 1:1000, Cell Signaling), rabbit-anti-GFP (polyclonal, 1:2000, #ab290, Abcam), rabbit-anti-copGFP (polyclonal, #PA5-22688, 1:5000, Invitrogen), rabbit-anti-Myc (monoclonal, #71D10, 1:1000, Cell Signaling), anti-mouse and anti-rabbit HRP secondary antibodies (1:10000, #A4416 and #A6154, Sigma). The membranes were visualized by ECL kits (#NEL105001EA, PerkinElmer; #34580, Thermo Scientific).

### 2.4. RNA extraction, reverse transcription (RT), quantitative real-time PCR (qPCR) and RNase R treatment

Total RNA was extracted by RNA-easy Isolation Reagent (#R701-01, Vazyme) according to manufacturer’s protocol. cDNA was generated by GoScript kit (# TM316, Promega) using 4 μg total RNA with random primers. qPCR was performed with Bio-Rad CFX96 using SYBR Master Mix (#Q311-02, Vazyme). Relative expression was calculated by the ΔΔCT method. 1μg total RNA was incubated with 4U/μg RNase R (#RNR07250, Lucigen) in 10μl reaction system at 37°C for 20min. Then the reverse transcription was performed immediately. RNase R was replaced by water in mock reactions. Primer information is provided in Table S2.

### 2.5. DNA extraction and Sanger sequencing

The PCR products were purified from DNA gels by extraction kit (#20051, Qiagen), and were directly sent for Sanger sequencing with gene-specific primers by Sangon Biotech.

### 2.6. Immunoprecipitation assay

Anti-FLAG M2 beads (#M8823, Sigma) were used according to the technical bulletin. Specifically, 20μl precleared M2 beads were added into 80μl cell lysate (120μl lysis buffer per well for 6-well plate), and the final volume was brought to 500μl by adding TBS. The sample was incubated at 4°C for 3h, washed with 500μl TBS for 3 times, and eluted by 10μl 2X SDS-PAGE buffer for Western blot analysis.

## 3. Results

### 3.1. Circular GFP RNA can be translated into functional protein

Previous research has reported that overexpressed circular GFP RNA without other modifications can be translated [9]. To test whether the ZKSCAN1 minigene circRNA overexpression vector is feasible for translatable circRNA study, split GFP was engineered into the vector and transfected into HEK293T cells. Robust GFP protein synthesis was detected by anti-GFP Western blot, but no fluorescence observed probably due to the protein misfolding of GFP tandem (data not shown). Then we inserted a P2A self-cleavage peptide sequence between two split GFP fragments (circ-GFP), which enables ribosome skipping and cleavage of the nascent polypeptide but without translation termination (Fig. S1A) [19,31]. Functional GFP protein translated from circ-GFP was detected by both fluorescence microscopy (Fig. S1B) and anti-GFP Western blot (Fig. S1C). Compared with the linear counterpart (lin-GFP), circ-GFP showed less translation efficiency but more resistance to co-expressed 4E-BP1, a cap-dependent translation inhibitor [7,32]. Consistent with the reported study [9], overexpressed circular GFP RNA can be translated cap-independently into functional protein.

### 3.2. Pervasive translation signals from overexpressed circRNAs

We next evaluated the coding potential of 30 human circRNAs (Table 1). (1) AUG circRNAs: the circRNAs are derived from exon circularization including the annotated AUG start codons. We expect AUG circRNAs have more tendency to initiate translation, although this still attracts controversy [15]. (2) Infinite ORF: no stop codon in their circular annotated ORF.

We cloned circRNA sequence into ZKSCAN1 minigene vector, and fused FLAG tag immediately downstream of the splice acceptor so that only when circRNAs are formed and translated can FLAG-fusion proteins be visualized by anti-FLAG Western blot analysis (Fig. 1A). No AUG start codon is located before FLAG in the linear ORF. The NeoR/KanR site in pcDNA3.1 vector was replaced by copGFP for estimating transfection efficiency and qPCR normalization. To validate the synthesis and the circularization of overexpressed circRNA, divergent FLAG and gene-specific primers were designed to perform RT–PCR (Fig. 1B) and BSJ sequencing (Table S1). As expected, ZKSCAN1 minigene vector accurately facilitates the production of most circRNAs except for circ-HSPB11 and circ-TNRC6B. Surprisingly, pervasive translation signals were obtained from more than 10 circRNAs by anti-FLAG immunoblotting (Fig. 1C). Circ-BBX (414nt), abbreviated as circ-BBX, shares two exons with circ-BBX (678nt), and both BBX circRNAs generated translation signals, indicating similar initiation mechanism underlies their translation. Circ-SYMPK and circ-ZFAND6 were reported to associate with ribosomes in ribosome profiling [33], and we indeed observed flagged protein produced from them. More translation signals will be captured if transfection efficiency is pushed highly enough (data not shown). Our further study focused on circ-BBX, circ-MAD1L1, circ-PPP1R13L, circ-SYMPK and circ-ZFAND6, for their particularly strong signals. Their circularity was evidenced by RNase R treatment as linear RNAs were digested while circRNAs not (Fig. 1D).

**Figure 1.**
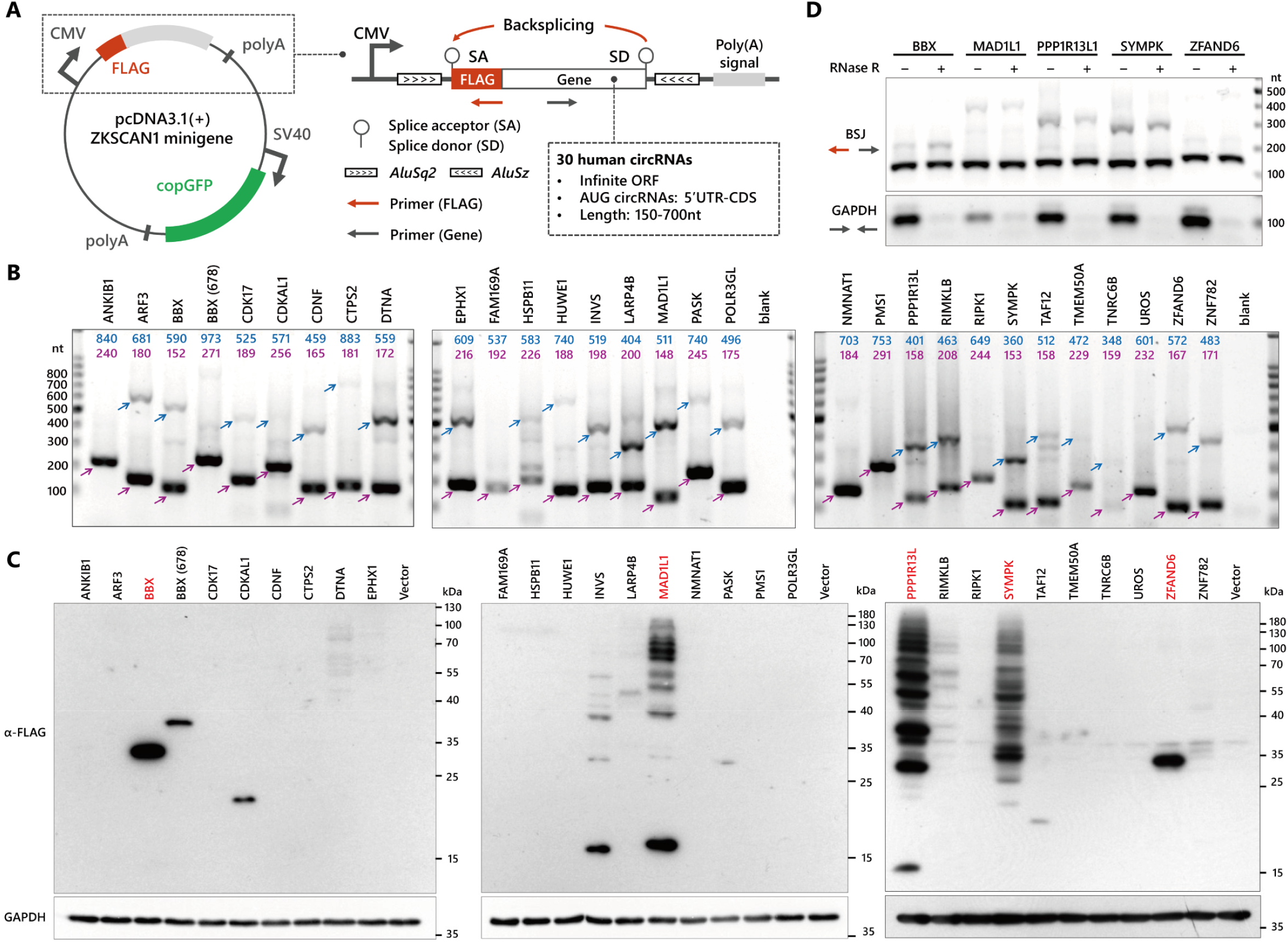
Pervasive translation events detected from overexpressed circRNAs. A. Schematic illustration of circRNA overexpression vector for coding-potential assessment. B. The rolling circle RT-PCR products from overexpressed circRNAs with divergent FLAG and gene-specific primers. The purple/blue arrows and numbers denote the 0-cycle/1-cycle RT-PCR products and their theoretical length. C. Western blot analysis with anti-FLAG antibody on proteins from HEK293T cells transfected with FLAG tagged circRNA overexpression plasmids. GAPDH as loading control. Five circRNAs for further investigation are marked in red. D. RNase R treatment for verifying the circularity of overexpressed circRNAs. Divergent FLAG and gene-specific primers were used for detecting circRNAs. GAPDH as linear RNA control.

### 3.3. Mutagenesis identifies aberrant translation from overexpressed circRNAs

We designed 3 subsets of mutants to examine whether the flagged proteins are translated from circRNAs (Fig. 2A). Firstly, the deletion of the upstream *AluSq2* sequence impaired, but not abolished, the formation of circRNAs, validated by RT-PCR and qPCR (Fig. 2D, 2G). This observation is consistent to former study [6]. Secondly, we mutated the splice acceptor (AG to AC) to create SA-mut constructs. For SA-mut, the genesis of circ-PPP1R13L and circ-MAD1L1 was abrogated, but other 3 circRNAs formed new BSJ using other splicing sites on vector, which were confirmed by RT-PCR (Fig. 2D) coupled BSJ sequencing (Fig. S2A). Thirdly, we asked whether cryptic translation initiation site is formed from cryptically spliced linear RNAs and promotes flagged protein translation. A stop codon was inserted at the end of circularized exon to test it. The circRNA-derived proteins will be masked in anti-FLAG Western blot as the translation terminates just before FLAG tag, but linear RNA-derived proteins can be visualized (Fig. 2B). As expected, the flagged protein synthesis was abolished in all SA-mut groups (Fig. 2C). Most ladder-like rolling circle translation signals disappeared in circ-MAD1L1-ΔALU, circ-PPP1R13L-ΔALU and circ-SYMPK-ΔALU. However, for all the ΔALU mutants, the flagged proteins were yielded when the genesis of circRNAs was curbed. The FLAG signals were still observed in circ-BBX-stop and circ-ZFAND6-stop, and their molecular weight was close to the wild-type (WT). These results demonstrate the flagged proteins in circ-BBX and circ-ZFAND6 groups are not translated from circRNAs. In a more comprehensive analysis of circ-PPP1R13L (Fig. 2E, 2F), the 15kD protein band remained upon splice donor (GT) removed or downstream *AluSz* sequence deleted, which hindered backsplicing (Fig. 4G, 4H). It rather corroborates the conclusion that the peptide of 15kD is not generated from circ-PPP1R13L. We also infer from these data that the expected splice acceptor, but not the assumed splice donor, is required for the genesis of some translation noise. There is a discrepancy between this observation and previous findings [6,28]. With stop codon inserted immediately behind FLAG (F+stop) or frame shifting (UTR+1, CDS+1) (Fig. 2E), rolling circle translation was disrupted, but the translation noise differs (Fig. 2F). So, on the plus side, substantial *Alu*-dependent flagged rolling circle translation may originate from circRNAs; on the minus side, concurrent *Alu*-independent noise obfuscates *bona fide* circRNA translation signals. Remarkably, we mutated all 5 annotated AUG codons to AUC on circ-PPP1R13L but failed to perturb the translation pattern (AUG-mut, Fig. 2F), indicating the translation of circ-PPP1R13L may be non-AUG mediated.

**Figure 2.**
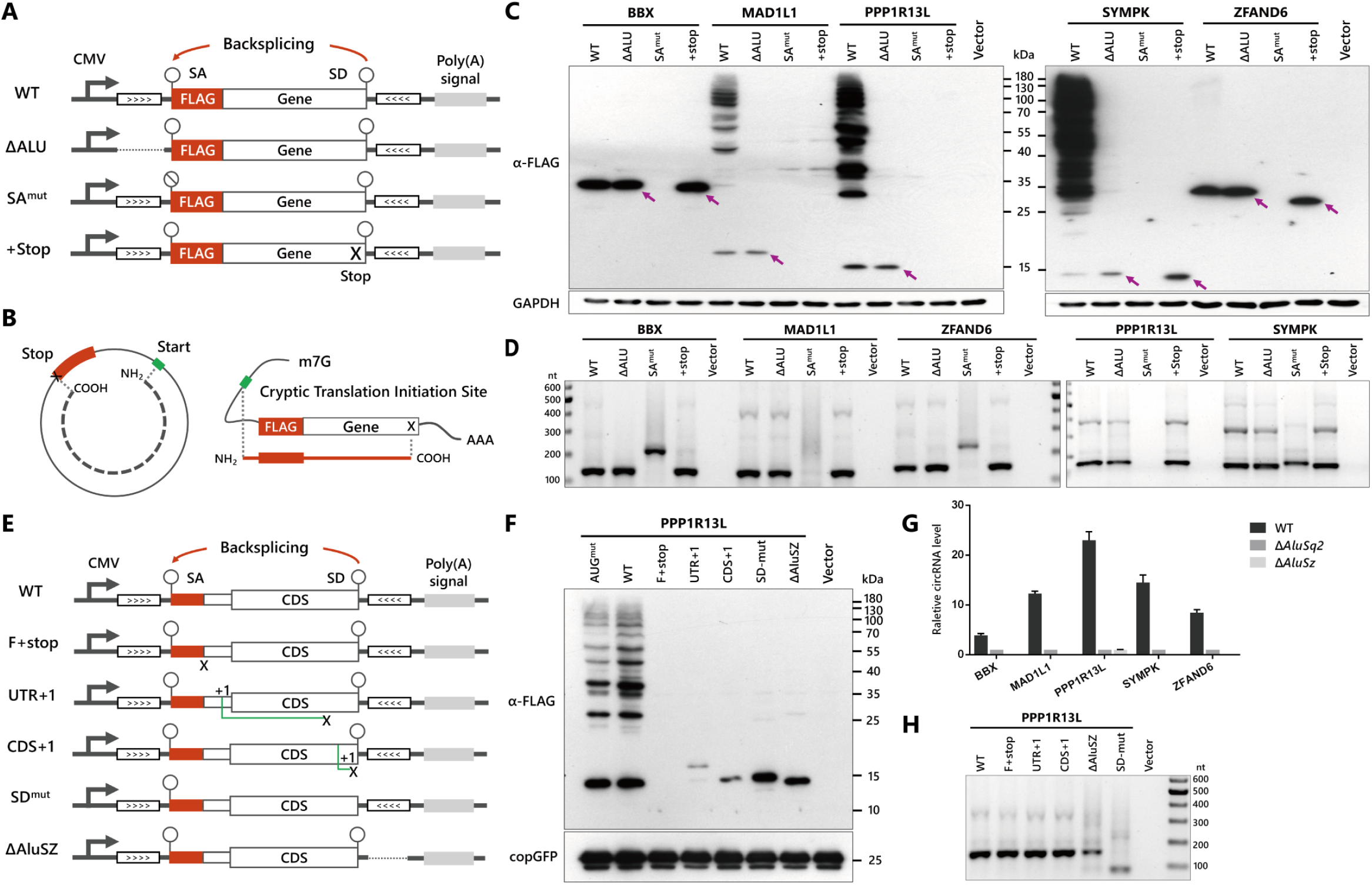
Mutagenesis-based validation of the translation from overexpressed circRNAs. A. Scheme overview of mutants for translation validation. Specifically, ΔALU: the upstream *AluSq2* sequence was deleted; SA-mut: the splice acceptor AG was mutated into AC; +Stop: a stop codon was added at the end of circRNA sequence. B. Scheme showing +stop mutants enable identification of the circRNA translation from linear RNA output. C and F. Western blot for translation validation. The protein bands regarded as translation noise are marked by arrows. GAPDH or copGFP as loading control. D and H. RT-PCR assay for the overexpressed circRNAs and their mutants with divergent FLAG and gene-specific primers. E. Scheme overview of circ-PPP1R13L mutants. Briefly, F+stop: a stop codon was inserted immediately downstream of FLAG; UTR+1: a cytosine was inserted into the untranslated region (UTR); CDS+1: a cytosine was added near the end of the coding sequence (CDS); SD-mut: the splice donor AG was removed; ΔAluSz: the downstream *AluSz* sequence was deleted. New ORFs created by frame shifting are shown as green lines. G. qPCR for the measurement of relative circRNA level using divergent FLAG and genes-specific primer sets. CircRNA expression was normalized to the ΔAluSq2 groups, and copGFP was used for calibration. Data is presented as mean ±SD (n=3).

### 3.4. Non-AUG start codons may initiate the translation of circRNAs

To further investigate the translation initiation of circRNAs, we mutated all the annotated AUG into GGG codons which are acknowledged not to initiate translation (Fig. 3A, 3B) [34]. The rolling circle translation of circ-MAD1L1 was significantly impeded by AUG mutation, but the aberrant translation remained. AUG mutation did not affect the production of circ-MAD1L1 (Fig. S2B). The translation noise of circ-BBX and circ-ZFAND6 stayed largely unaffected in AUG mutants, suggesting the aberrant translation is initiated by the elements beyond circRNA sequence, a scenario distinct from circ-ZNF609 [28,29]. The AUG mutation was also unable to abrogate the rolling circle translation of circ-SYMPK (Fig. 3B). With the same method, we tested circ-PAPOLA, a circRNA that associates with ribosome [33], carries infinite ORF, but naturally lacks annotated AUG codon (Table. 1). The rolling circle translation of circ-PAPOLA were indeed observed (Fig. 3B). These results indicate the non-AUG start codons may initiate the translation of circRNAs.

**Figure 3.**
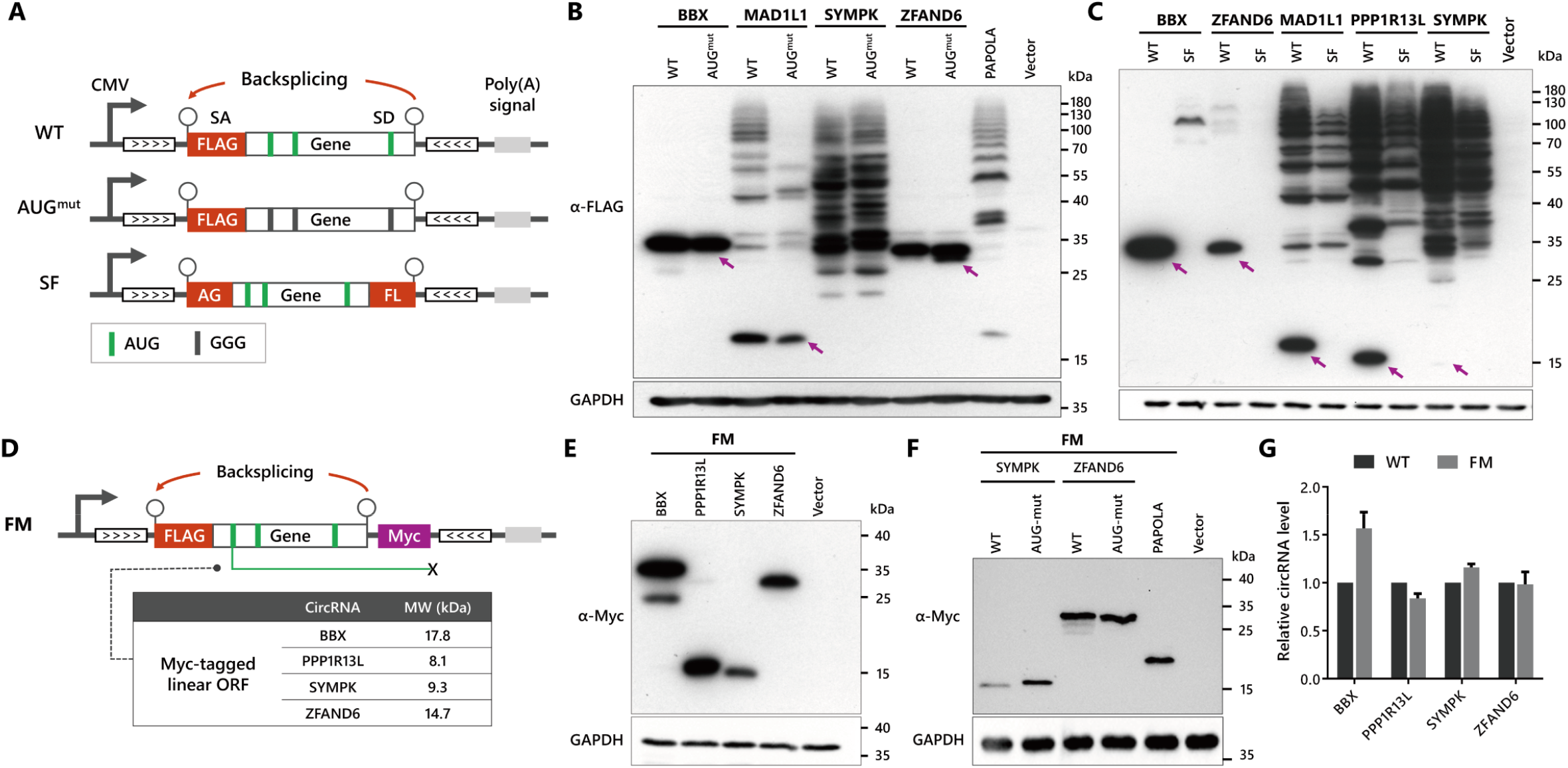
Analysis of the aberrant translation from overexpressed circRNAs. A. Scheme overview of constructs. AUG-mut: all the annotated AUG codons were mutated into GGG; SF: FLAG tag was split into two parts flanking circRNA sequence. B and C. Western blot analysis of HEK293T transfected with AUG-to-GGG mutant and split FLAG constructs. GAPDH as loading control. The fallacious translation signals are marked by arrows. D. Schematic representation of FLAG-Myc reporter design. Myc tag was embedded into the end of linear ORF beyond backsplicing cassette. The putative Myc tagged ORF is presented as green line, and the molecular weight information is listed. E and F. Western blot of HEK293T transfected with FM and the derivative mutants. All the annotated AUG start codons were mutated into GGG codons in AUG-mut constructs. GAPDH as loading control. G. CircRNA levels of WT and FM were measured by qPCR with divergent FLAG and gene-specific primers. CircRNA expression was normalized to WT. GAPDH was used for calibration. Data is presented as mean ±SD (n=3)

### 3.5. The aberrant translation is arisen from coexistent linear RNAs

To disentangle the authentic translation signals from noise, we flanked circRNA sequence with split FLAG to create SF constructs (Fig. 3A). Since the previously verified fallacious signals disappeared, it seems to be an efficacious noise masking strategy (Fig. 3C). Strangely, we detected new protein bands (∼100kD) in circ-BBX-SF, which were never seen in WT. More flagged proteins (∼100kD) in circ-ZFAND6 were occasionally visualized depending on transfection efficiency. Inspired by the observation of circ-PPP1R13L-SD-mut (Fig. 2E), we hypothesized that the putative intronic splice donor is involved in the aberrant translation. FLAG-Myc dual tag reporter (FM) was designed to test it. Myc tag was inserted at the end of linear ORF between splice donor and *AluSz* sequence, which is not supposed to be backspliced (Fig. 3D). Therefore, a part of translation noise from linear RNAs can be labeled by the anti-Myc immunoblotting signals. Such engineering did not inhibit the production of overexpressed circRNAs (Fig. 3G), a finding that aligns with recent study [20]. For FM, anti-Myc signals were obtained, but their molecular weight was larger than the theoretical value of putative linear ORF (Fig. 3D, 3E). Moreover, AUG mutation was also unable to disturb the synthesis of Myc tagged proteins (Fig. 3F).

Considering flagged and Myc-tagged noise shows similar SDS-PAGE gel shift and resists AUG mutation, we asked whether they are from the same proteins. We further performed immunoprecipitation (IP) to purify FLAG fusion proteins from cell lysate and visualized them by anti-Myc Western blot (Fig. 4A). Strong Myc signals representing previously defined aberrant translation were detected in FM IP groups, but not in SFM IP groups in which FLAG is separated (purple arrows in Fig. 4B). By virtue of IP enrichment, robust anti-Myc rolling circle translation signals unexpectedly came out. Moreover, translation noise was even observed in SFM IP groups of circ-PPP1R13L and circ-SYMPK (blue arrows in Fig. 4B), indicating linear RNAs involving intact FLAG BSJ region were created and translated. We excluded the possibility of novel circRNAs containing both FLAG and Myc sequence by RT-PCR using corresponding divergent primers (Fig. 4A, 4C). Eventually, we concluded that the heterogeneous aberrant translation originates from concomitant linear RNAs, and directly demonstrated that the circRNA-iconic rolling circle translation signals can be stimulated by linear transcripts, most likely by RNA concatemers (Fig. 4D).

**Figure 4.**
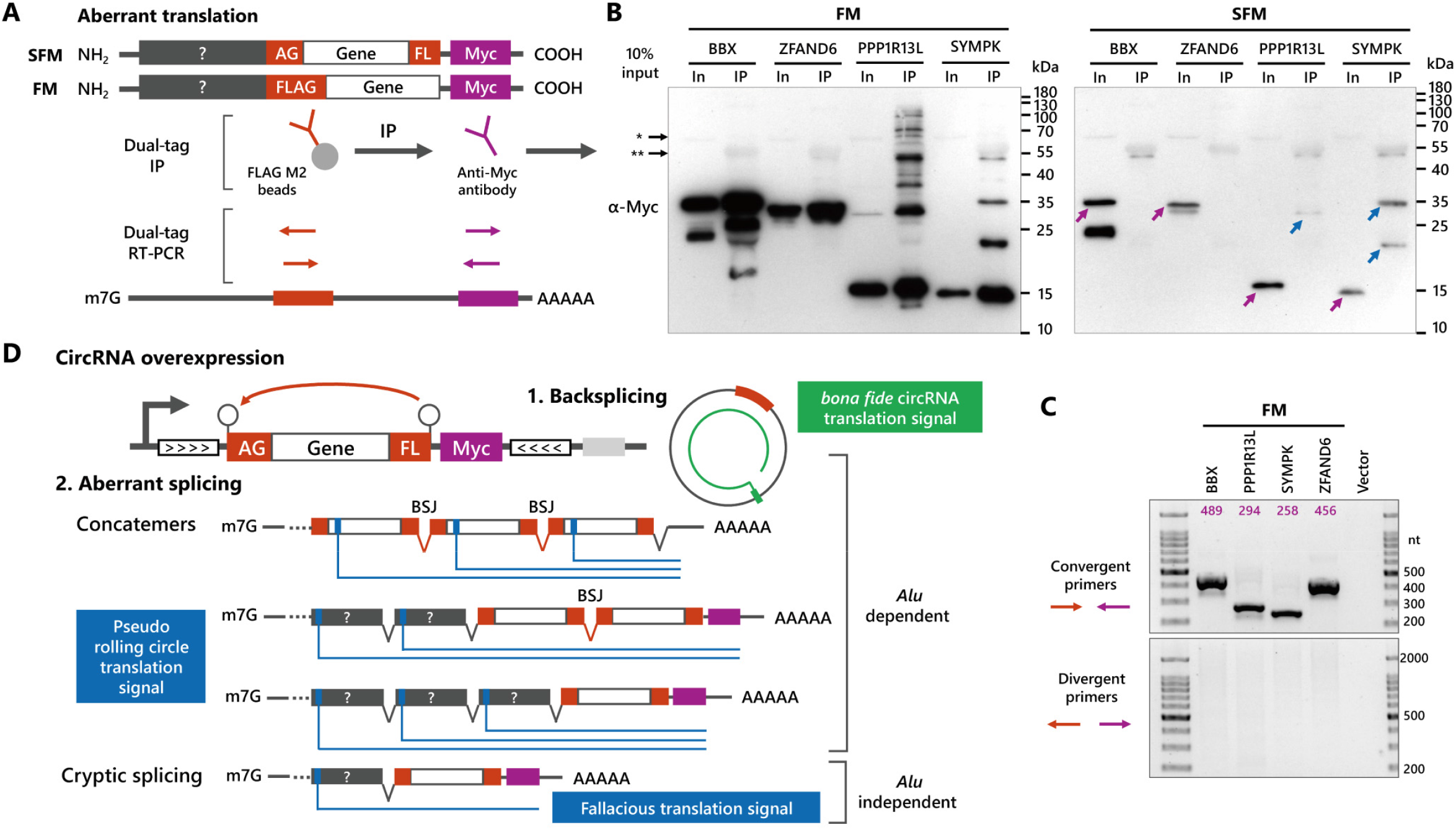
Heterogeneous outputs from complementary intron-mediated circRNA overexpression vector. A. Workflow for the examination of aberrant translation by dual-tag IP and RT-PCR. B. FLAG IP coupled anti-Myc Western blot analysis of HEK293T transfected with FM and SFM constructs. Blue/purple arrows represent fallacious translation signals from linear transcripts with/without BSJ. The single and double asterisks denote the non-specific antibody binding to endogenous proteins and FLAG antibody fragments, respectively. C. RT-PCR analysis of HEK293T transfected with FM constructs with divergent primers of FLAG and Myc to detect possible circRNAs involving both FLAG and Myc sequence. RT-PCR using convergent primers acts as control group, and the theoretical length of PCR products was denoted. D. Proposed heterogeneous outcomes of circRNA overexpression. The authentic translation from circRNAs is described as green curve, and the linear RNA derived translation is illustrated as blue lines.

## 4. Discussion

Translate or not? Our endeavors, technically, have not completely confirmed that the rolling circle translation are derived from circRNAs in this study, because BSJ-bridging epitope tag and mutagenesis done here cannot circumvent the latent artefacts arisen from RNA concatemers or other unforeseen transcripts (Fig. 4D). These byproducts, which share the same BSJ region with circRNAs, escape from most PCR-based discrimination and can be only visualized via Northern blot. Even worse, the rationale evidencing translatable circRNAs usually rests on three pillars: immunoblotting to detect backsplicing-specific proteins, mass spectrometry to determine BSJ-specific peptides, ribosome profiling to identify ribosome-protected BSJ sequence; coexistent linear concatemers including BSJ will take a sledgehammer to all of them if conclusions heavily rely on overexpression without careful control experiments. A discernable connection between translation signal and flanking *Alu* sequence is a prerequisite but not sufficient condition to prove circRNA translation because the complementary *Alu* sequence could also contribute to the formation of undesired linear transcripts and exon concatemerization. Exemplified by our study, the practical limitations for detecting and avoiding unwanted transcripts would ambush the overexpression-based circRNA investigation. Fortunately, the caveat echoes, the remedy follows [3,4,14,16,35]. Given most of RNA concatemers are observed in complementary intron-mediated backsplicing, other rarely used circRNA overexpression methodologies may overcome this dilemma (such as the strategy of inserting Quaking binding sites [18,36] or using twister-optimized RNA expression [37]) and deserve a further evaluation.

Overexpression of circRNA carrying infinite ORF, resulting in a plethora of unfolded repeating polypeptides, can overwhelm the cellular protein quality control (QC) system to artificially amplify the signals of naturally degraded proteins, supported by the recent finding that rolling circle translation products undergo proteasome-mediated degradation[9]. However, too much attention has been attracted on justifying the existence of circRNA-derived protein to realize translation outcome is a total package enveloping entire translation process, which could shed the enigma of ribosome-associated but peptide-undetected circRNAs. It is very tempting to speculate translatable circular RNAs function not by the intact translated proteins, but by the translation process itself, to regulate cellular activities. Because once a circRNA is translating, it must be strictly and precisely governed by translation machinery and QC system, of which dysregulation is sufficient to raise functional impact. Impressive gains in the understanding of ribosome quality control [38] invite our further hypothesis: unstopped circular ORF renders ribosome collision an imminent possibility, because translating circRNAs threading through ribosomes without stop codon-mediated termination are prone to becoming fully loaded and susceptible to ribosome stalling, whereas the relevant research stays rather scarce.

Finally, non-AUG start codons have emerged as important translation initiators with profound cellular function [39,40]. Although our results regarding non-AUG translation could be preliminary, we speculated that non-AUG may function more efficiently in triggering rolling circle translation of circRNAs, compared with the scenario in linear mRNA. Non-AUG circRNA translation has been also observed by another group with more solid mutagenesis evidence [9].

The optimism about endogenous translatable circRNAs would be tempered if flawed experiments piggyback leapfrogging functional studies. More interrogation and development of circRNA research toolset merits spotlight in the future.

## Acknowledgments

We thank Miaowei Mao and Tao Zu for the advice about *in silico* splicing signal prediction and non-AUG translation. This work was supported by National Natural Science Foundation of China (81874249), Guangdong Basic and Applied Basic Research Foundation (2020A1515011125), and Shenzhen Basic Research Grants (JCYJ20180223181224405, JCYJ20180507182657867).

## Author Contributions

Y.J. conceived the project, designed the experiments, performed the analysis, and drafted the manuscript. X.C. assisted manuscript editing. W.Z. supervised the project. W.Z. and X.C. acquired the funding.

## Competing interests

The authors declare no competing interests.

## Supplementary materials

**Figure S1.**
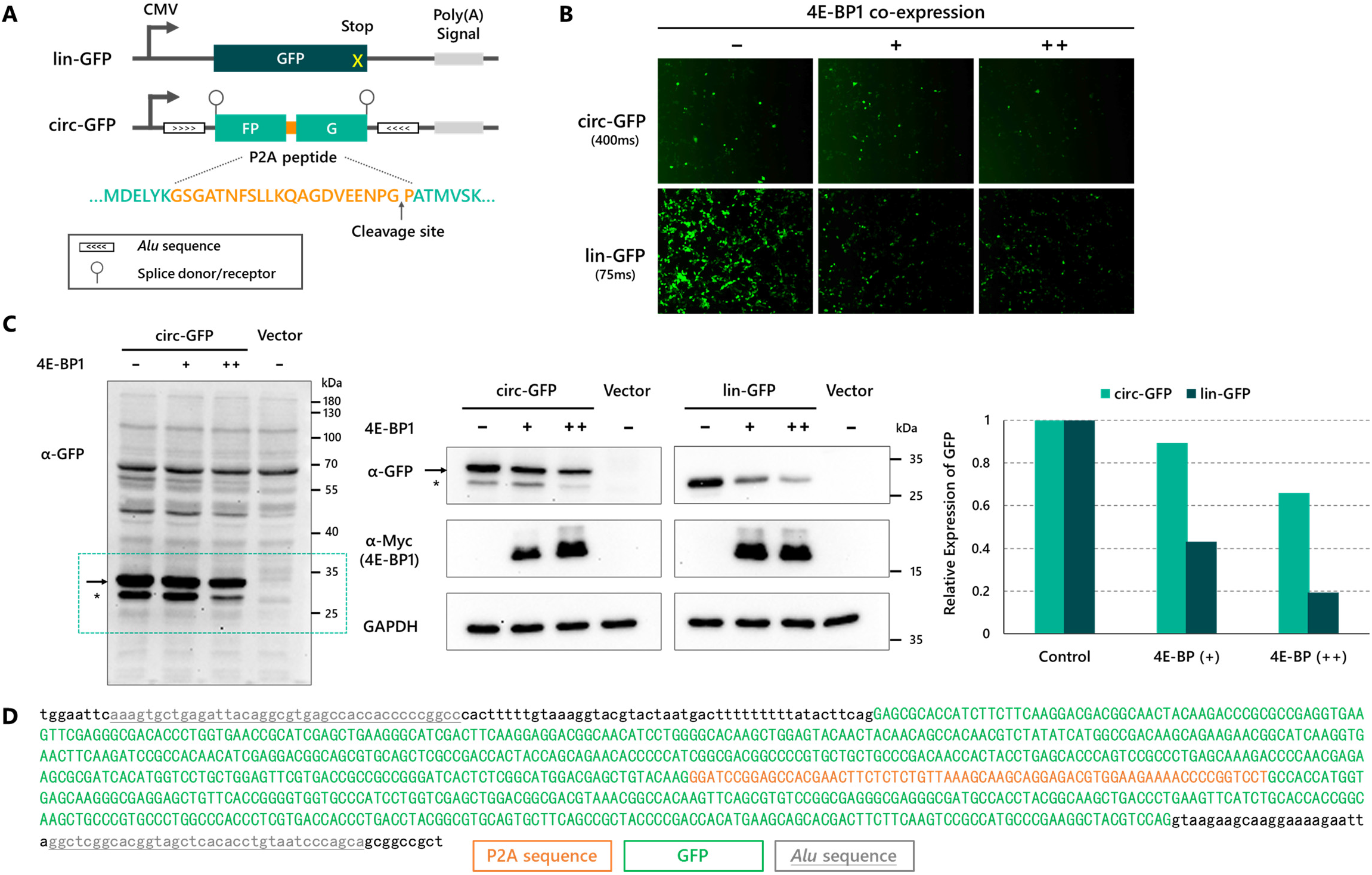
The functional translation of circular GFP RNA. A. Schematic diagram illustrating construct design of circular and linear GFP overexpression. B. Functional GFP proteins expressed by the circ-GFP overexpression are resistant to the activity of 4E-BP1. HEK293T cells were transfected with 1.5ng/well plasmids of circ-GFP or lin-GFP in 6-well plate, and 0, 0.5, 1ng/well Myc-4E-BP1, and 1, 0.5, 0ng/well control vector was used for co-transfection (totally 2.5ng DNA/well). After 40h, the fluorescence was observed and the exposure time was 400ms and 75ms for circ-GFP and lin-GFP, respectively. C. Western blot analysis of the samples in Fig. S1B. Left: the overview of anti-GFP immunoblotting in longer exposure, and the GFP monomer for analysis was selected by green dotted box. The asterisk denotes GFP fragment. Middle: Western analysis of GFP monomer expression in suitable exposure. Blots were hybridized with GFP, Myc and GAPDH antibodies. Right: quantification of GFP levels relative to control group (n=1). D. The sequence information of circ-GFP. Intronic sequence is in lower case.

**Figure S2.**
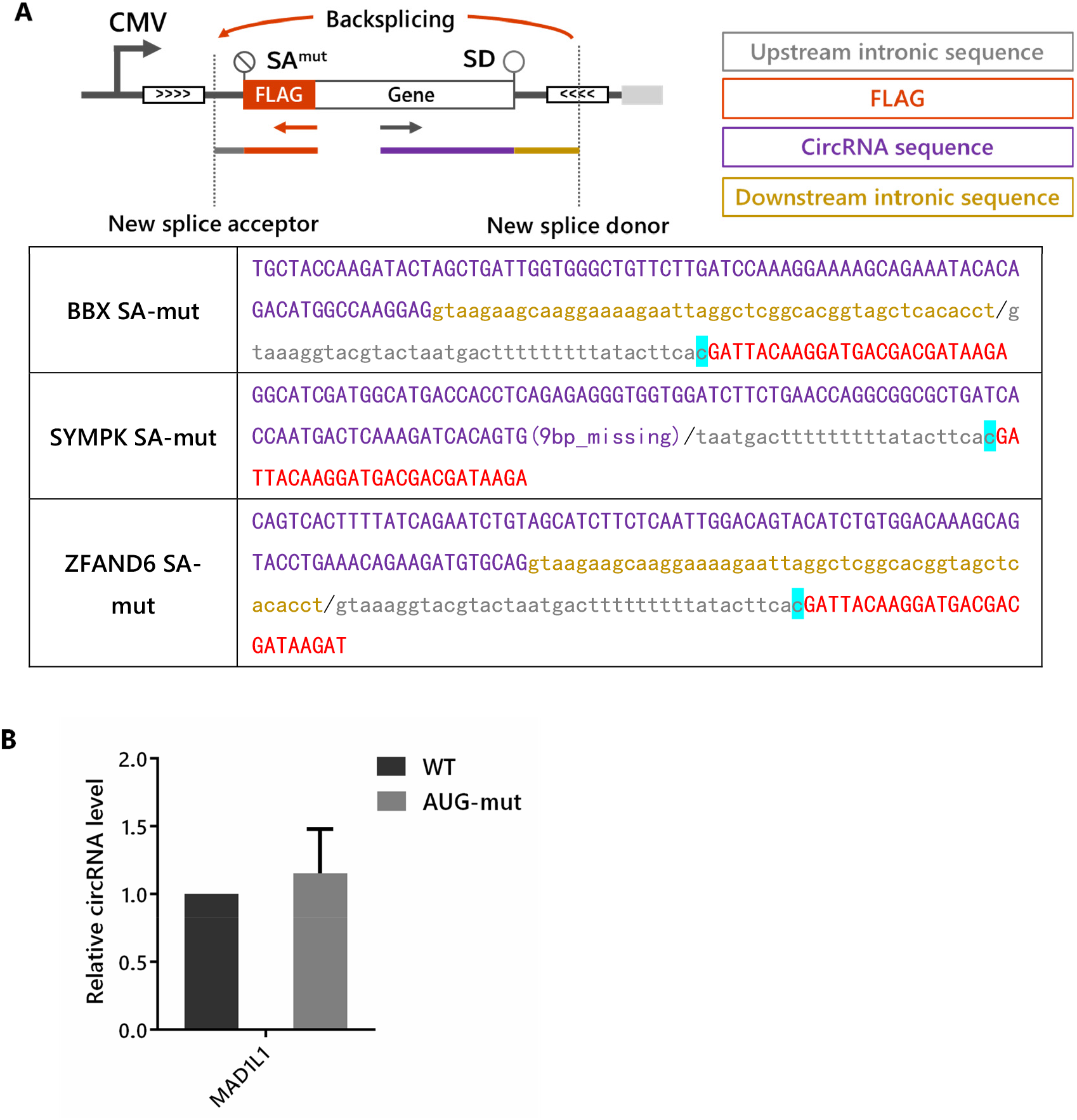
Analysis of the transcripts from AUG and splice acceptor mutants (related to Fig. 2D and 3B) A. Sequencing analysis of new BSJ formed from splice acceptor mutants. An illustration describing new backsplicing from other splicing sites on vector when the putative splice acceptor is mutated. Intronic sequence is in lower case, the mutated cytosine is highlighted in cyan, and the proposed new BSJ is presented as slash. B. CircRNA levels of MAD1L1 were measured by qPCR with divergent FLAG and gene-specific primers. CircRNA expression was normalized to WT. GAPDH was used for calibration.

**Figure S3.**
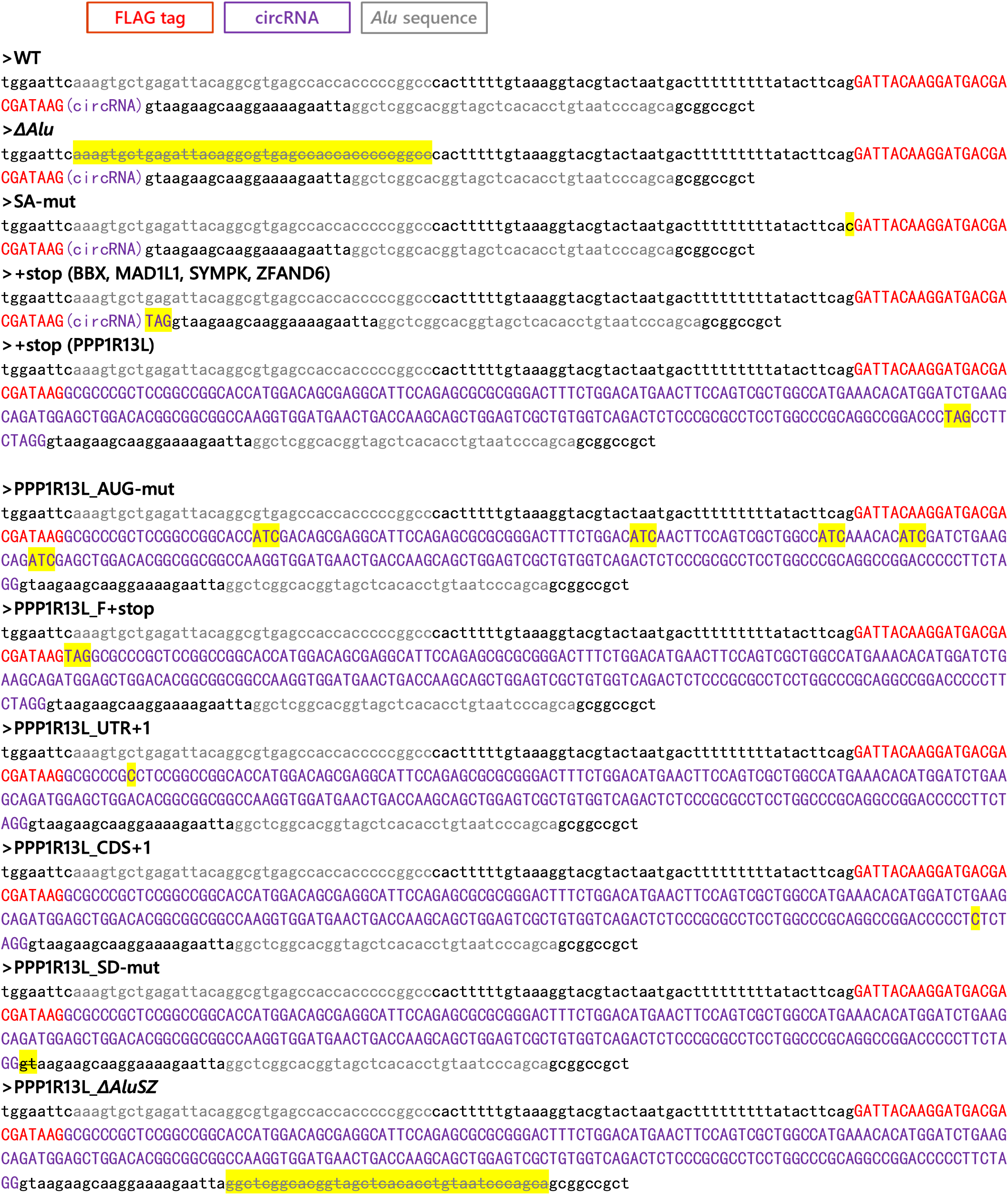
Sequence information of circRNA constructs (related to Fig. 2) The sequence of circRNA overexpression constructs used in this study. FLAG tag (red), circRNA sequence (purple) and *Alu* sequence (gray) are presented, intronic sequence is in lower case, and the mutation is highlighted in yellow.

**Figure S4.**
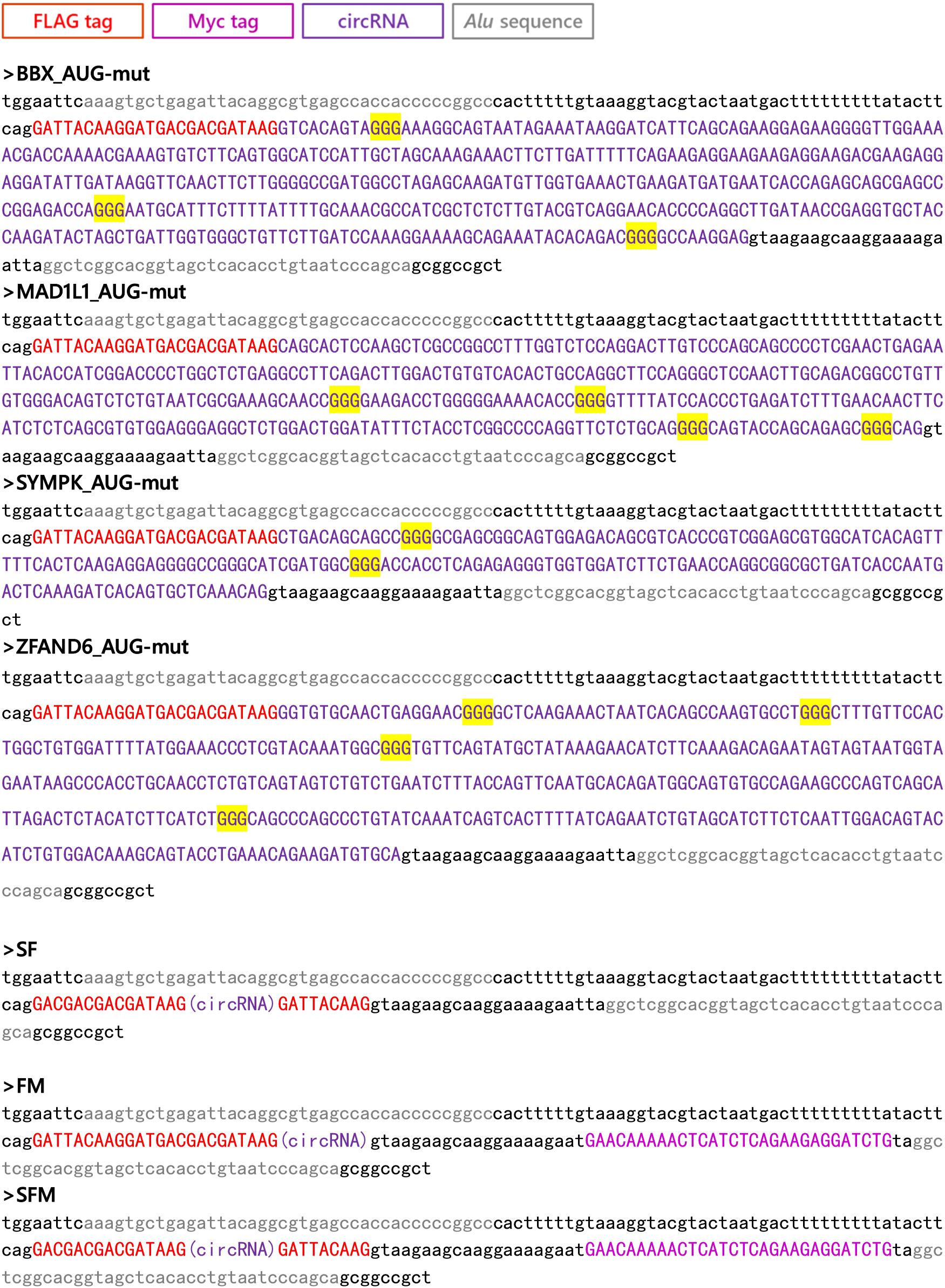
Sequence information of circRNA constructs (related to Fig. 3 and Fig. 4) The sequence of circRNA overexpression constructs used in this study. FLAG tag (red), Myc tag (magenta), circRNA sequence (purple) and *Alu* sequence (gray) are presented, intronic sequence is in lower case, and the mutation is highlighted in yellow.

**Table S1.**
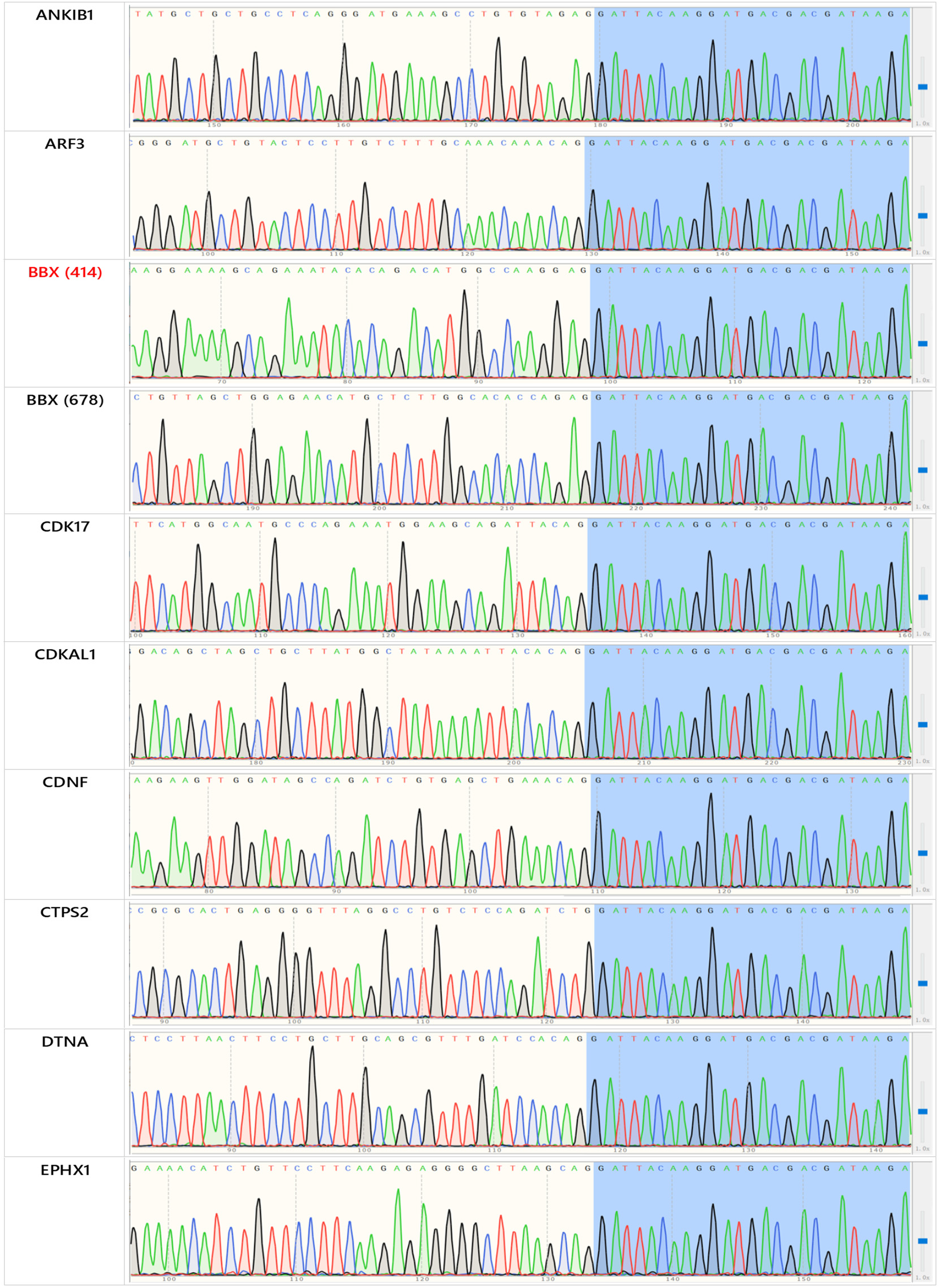
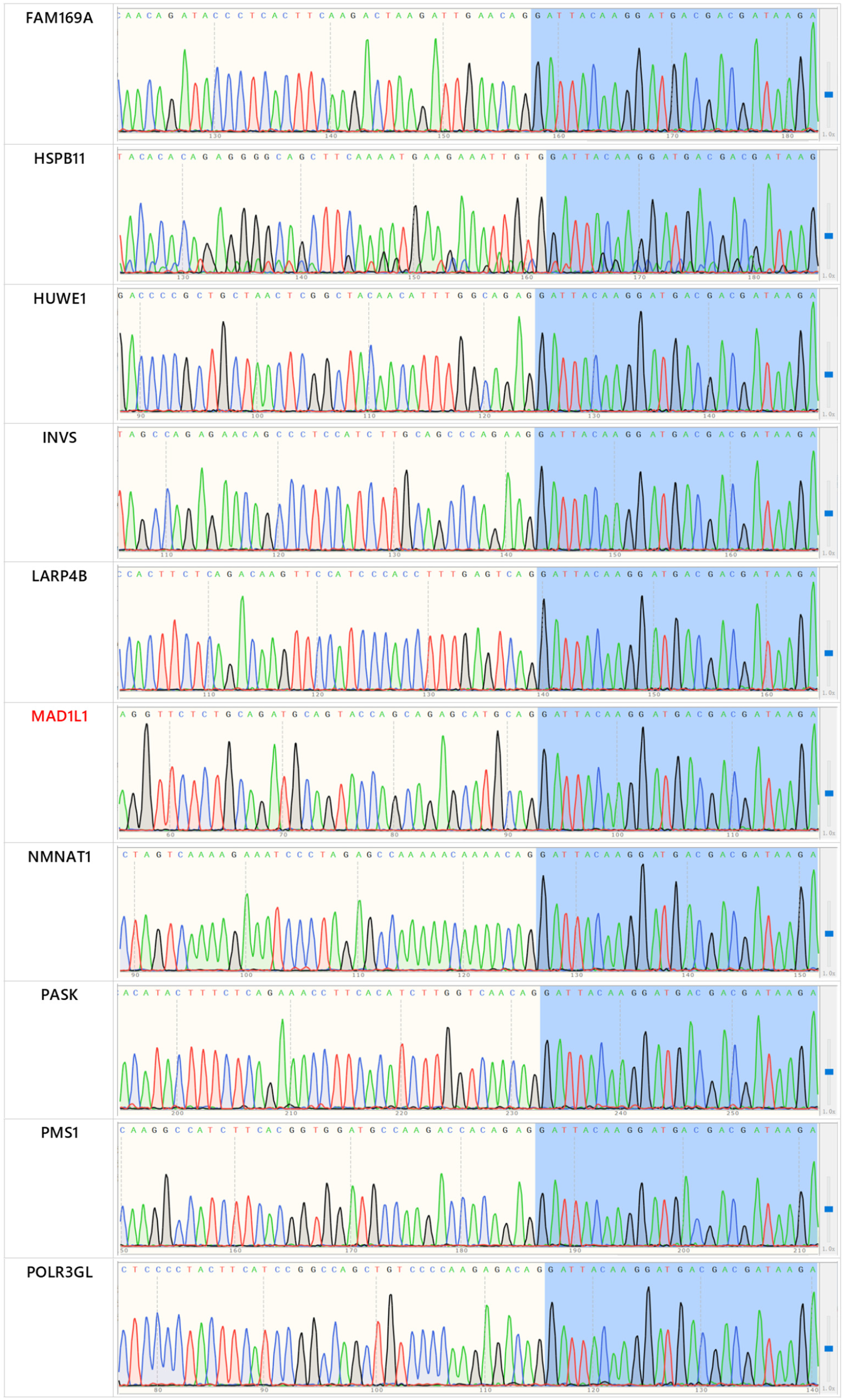
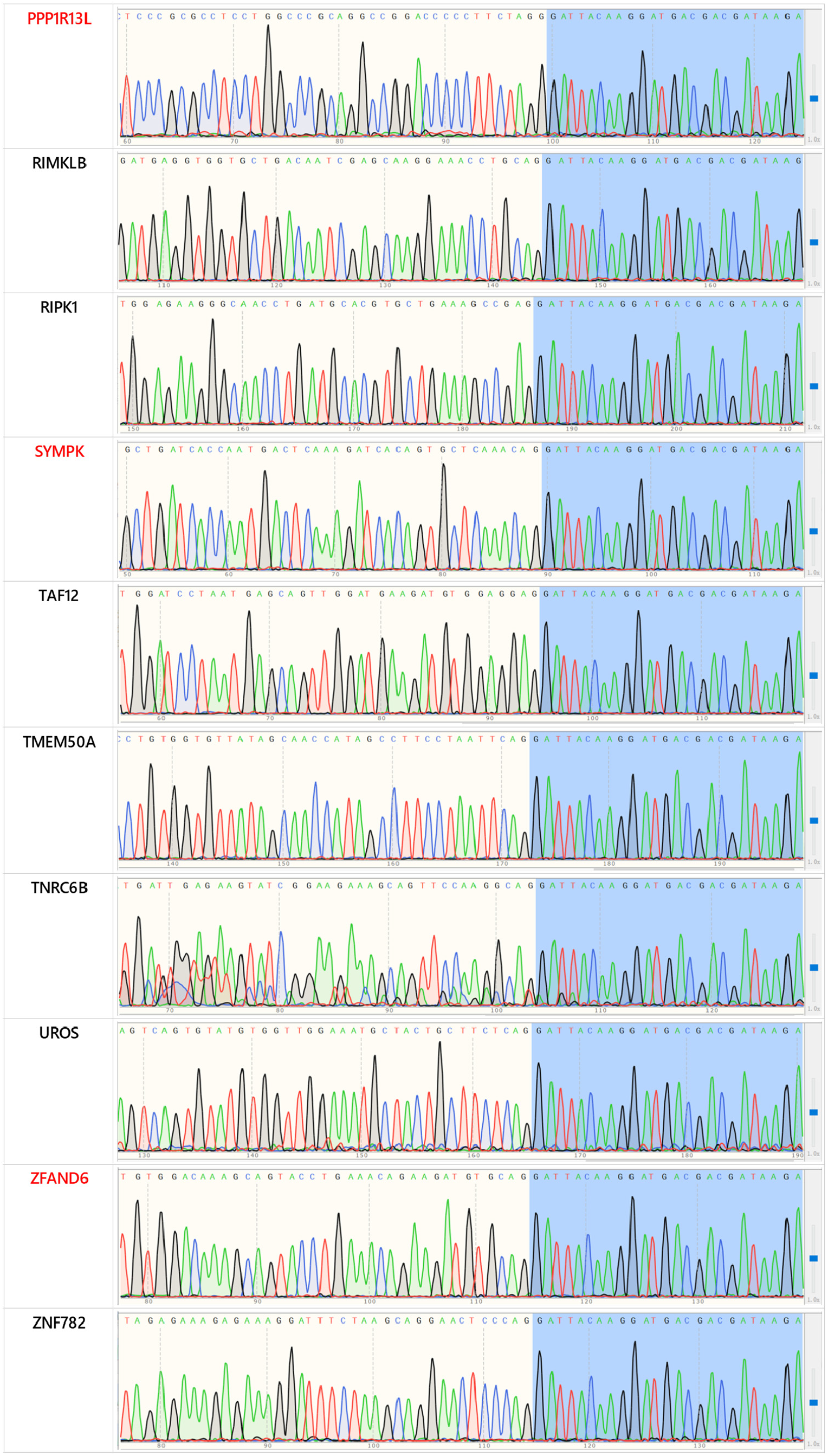
BSJ validation by Sanger sequencing. The FLAG sequence is marked in blue. The BSJ is represented by the junction of two background colors.

**Table S2.**
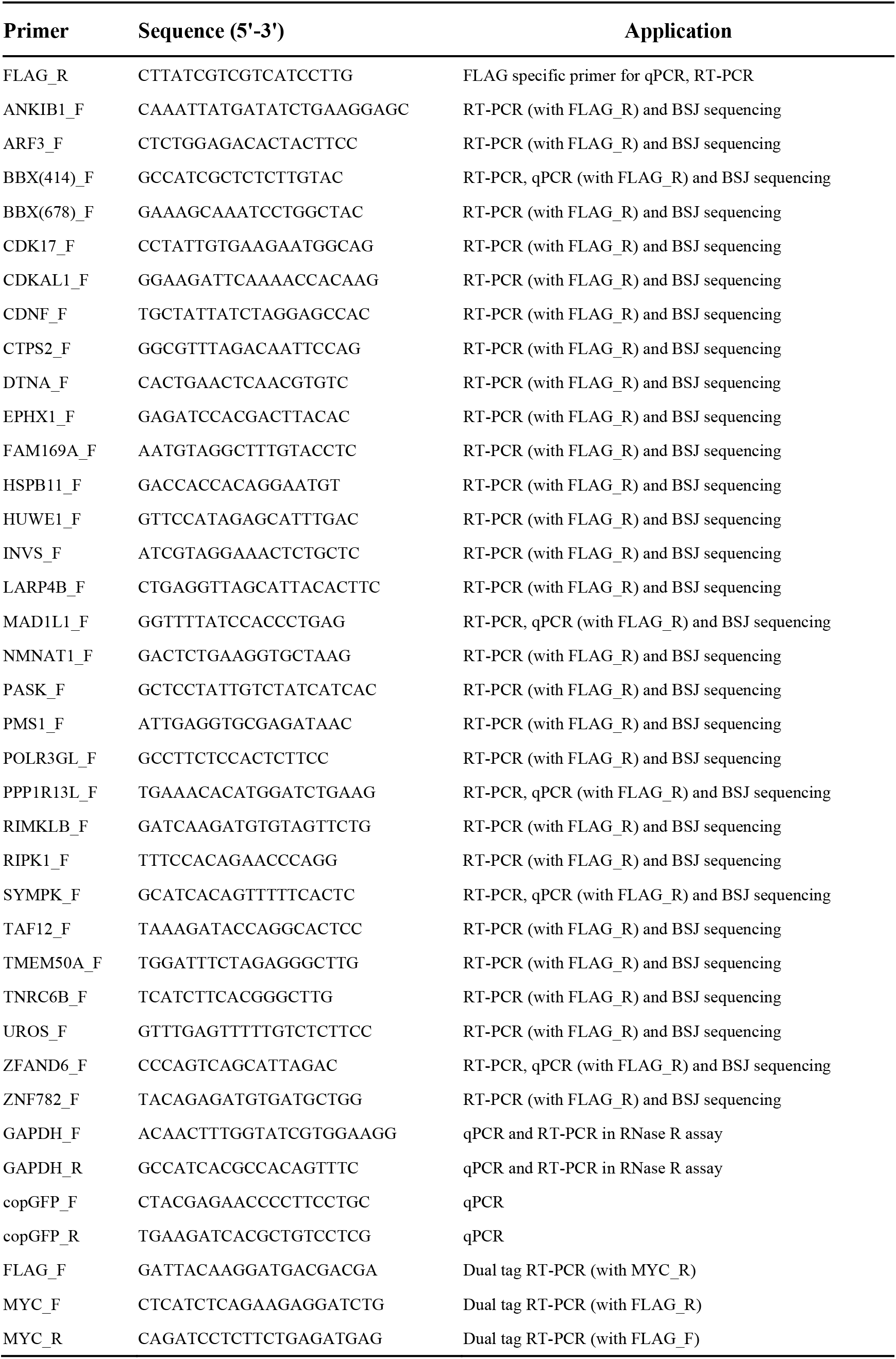
Sequence information of primers used in this study.

## Notes

### Competing Interest Statement

The authors have declared no competing interest.

